# Spontaneous mutations that confer resistance to 2-deoxyglucose act through Hxk2 and Snf1 pathways to regulate gene expression and HXT endocytosis

**DOI:** 10.1101/815118

**Authors:** Samantha R. Soncini, Dakshayini G. Chandrashekarappa, David A. Augustine, Kenny P. Callahan, Allyson F. O’Donnell, Martin C. Schmidt

## Abstract

Yeast and fast-growing human tumor cells share metabolic similarities in that both cells use fermentation of glucose for energy and both are highly sensitive to the glucose analog 2-deoxyglucose. Spontaneous mutations in *S. cerevisiae* that conferred resistance to 2-deoxyglucose were identified by whole genome sequencing. In addition to three aneuploid strains, we detected missense alleles of the *HXK2, REG1, GLC7* and *SNF1* genes that were shown to confer significant resistance to 2-deoxyglucose. All three missense alleles in *HXK2* resulted in significantly reduced catalytic activity. Missense alleles affecting the Snf1 kinase pathway (*REG1*, *GLC7* and *SNF1)* exhibited different capacities to affect the regulation of invertase expression. Of the seven missense alleles identified in this study, all but one affected hexose transporter endocytosis by increasing plasma membrane occupancy of the Hxt3 protein. Increased expression of the DOG (deoxyglucose) phosphatases has been associated with resistance to 2-deoxyglucose. Expression of both the *DOG1* and *DOG2* mRNA was elevated after treatment with 2-deoxyglucose. Deletion of the *HXK2* and *REG1* genes confers resistance to 2-deoxyglucose and causes increased expression of the *DOG2* mRNA. We conclude that Snf1 kinase-mediated regulation of the endocytosis of the hexose transporters and regulation of *DOG2* expression are important mechanisms for resistance to 2-deoxyglucose. However, the dominant *SNF1-G53R* allele can confer additional 2-deoxyglucose resistance in cells that are genetically compromised in both the endocytosis and *DOG* pathways. Thus at least one more mechanism for conferring resistance to this glucose analog remains to be discovered.

**Author Summary:** Yeast and fast-growing human tumor cells share metabolic similarities in that both cells use fermentation of glucose for energy and both are highly sensitive to the glucose analog 2-deoxyglucose. Another similarity between yeast cells and human tumor cells is that both cells can acquire resistance to 2-deoxyglucose, an outcome that can limit the usefulness of some cancer therapeutics. In this study, we used bakers’ yeast as a model organism to better understand the mechanism of toxicity and acquisition of resistance to 2-deoxyglucose. Spontaneous mutations in S. cerevisiae that conferred resistance to 2-deoxyglucose were isolated and identified by whole genome sequencing, a technology that was not available until recently. Our studies indicate that 2-deoxyglucose becomes toxic after it is phosphorylated by an enzyme called hexokinase. One important route to resistance is to reduce hexokinase activity. Other parallel pathways to resistance include increased expression of a hydrolase that degrades the toxic metabolite, altered localization of glucose transporters and altered glucose signal transduction pathways.

## Introduction

Genetic selection for mutations that confer resistance to the glucose analog 2-deoxyglucose (2DG) have been undertaken since the 1970s (1), and these studies have identified numerous genes that play roles in the maintenance of glucose repression (2–4). These screens were conducted on media that contained alternative carbon sources (e.g., sucrose, raffinose, galactose). Mutations that relieved glucose repression of gene expression were isolated and led to the hypothesis that 2DG promoted glucose repression, thus inhibiting growth on alternative carbon sources. However, recent studies with 2DG using glucose as the carbon source (5, 6) have challenged this hypothesis, since promotion of glucose repression should not by itself be toxic to cells growing on glucose. We discovered a link between the Snf1 signaling pathway, 2DG resistance, and the membrane localization of the high-capacity glucose transporters, Hxt1 and Hxt3 (7). Cells lacking Snf1 kinase are hypersensitive to 2DG (5) and have greatly decreased plasma membrane retention of Hxt1 and Hxt3 (7). This regulation of HXT localization is mediated by the *α*-arrestins Rod1 and Rog3, both of which are substrates of the Snf1 kinase (7, 8). An analogous signaling pathway is present in mammalian cells, where the AMP-activated protein kinase (AMPK), the mammalian ortholog of Snf1, regulates endocytosis of the glucose transporter GLUT1 by phosphorylating the TXNIP *α*-arrestin (9).

An alternative approach to studying resistance to 2DG was to screen for genes that conferred resistance when overexpressed from a high-copy plasmid. Such an over-expression screen identified the 2-deoxyglucose-6-phosphatate phosphatase genes, *DOG1* and *DOG2* (*10*). These genes encode phosphatases in the HAD-family (halo-acid dehalogenase) most closely related to the glycerol-3-phosphate phosphatases (*GPP1* and *GPP2*) with the signature catalytic motif DxDxT (11) present at their N-termini. A recent study of 2DG resistance using an unbiased proteomics approach found that increased expression of the DOG phosphatases was a critical response to the presence of 2DG and that multiple signaling pathways mediated the transcriptional regulation of the *DOG1* and *DOG2* genes (12). Furthermore, overexpression of a related human HAD phosphatase could impart 2DG resistance in human cell line, suggesting that this mechanism of resistance is conserved between yeast and human.

In this study, we undertook a new selection for 2DG-resistant strains using a few novel approaches. First, we selected for spontaneous 2DG-resistant strains growing on glucose rather than alternative carbon sources. Second, we included an HXT3-GFP reporter in our starting strains in order to determine if these mutations caused changes in the localization and membrane trafficking of this glucose transporter. Finally, we used whole genome sequencing to identify mutations in our resistant strains in an unbiased way. Using this approach, we defined new alleles in *HXK2* and *REG1* that confer 2DG resistance, demonstrate that both the *HXK2* and *SNF1* pathways regulate endocytosis and infer the existence of at least one additional pathway that can confer 2DG resistance.

## Results

### Isolation and identification of 2DG-resistant mutants

In order to obtain new, independent, and spontaneous 2DG-resistant mutants in *S. cerevisiae*, single colonies of wild-type yeast cells were grown overnight in synthetic complete media with 2% glucose as the carbon source. Cultures were then diluted in water, and 2×10^7^ cells were spread onto agar plates containing synthetic complete media with 2% glucose and 0.1% 2DG. Plates were incubated for 4-6 days until single colonies grew and could be streaked onto a fresh plate without 2DG. Isolation of 2DG-resistant strains using haploid cells usually generated 6-10 colonies per plate. Single colonies from these initial isolates were then tested for growth on media with and without 2DG. Since we showed previously that deletion of the *HXK2* and *REG1* genes conferred 2DG resistance to cells using glucose as the carbon source (5), we expected to recover loss of function alleles in these genes. Haploid strains were mated to *hxk2*Δ and *reg1*Δ strains, and the resulting diploids were scored for complementation. Twenty-eight 2DG-resistant haploid strains were examined. Of these, 21 were in the same complementation group as *hxk2*Δ, 1 was in the same complementation group as *reg1*Δ, and 6 were not in either of these complementation groups. Mating these 2DG-resistant strains with a wild type showed that 27 contained recessive alleles conferring 2DG resistance. One strain contained a dominant mutation.

In order to identify the mutations that confer 2DG resistance, we isolated genomic DNA from the 2DG-resistant strains and subjected it to whole genome sequencing using Illumina NextSeq500. This produced 151-bp paired-end reads (13) with an average depth of 20 million reads per sample. At this depth, each nucleotide in the yeast genome was sequenced an average of 240 times. We identified candidate mutations in 19 of the 22 haploid strains sequenced (Table 1). Of the many strains in the *hxk2*Δ complementation group, 15 were subjected to whole genome sequencing. Of these, 12 had the same mutation producing the missense allele, *hxk2-G55V*. We also isolated *hxk2-D417G* twice and *hxk2-R423T* once. We identified candidate mutations in four haploid strains that produced missense alleles in the *SNF1*, *REG1*, and *GLC7* genes. The one resistant strain with a dominant mutation contained the missense allele *SNF1-G53R.* This same allele had been isolated previously in a screen for Snf1 kinase function in the absence of the complex’s gamma subunit, Snf4 (14). Three recessive alleles produced missense mutations in the yeast protein phosphatase I complex comprised of Glc7 and Reg1 (11). We isolated *glc7-Q48K* twice and *reg1-P231L* once. Three strains that were 2DG resistant but lacked candidate mutations were found to be aneuploid with extra copies of multiple chromosomes (Fig S1). Further characterization of the aneuploid strains will be described in a subsequent publication.

**Table 1.**
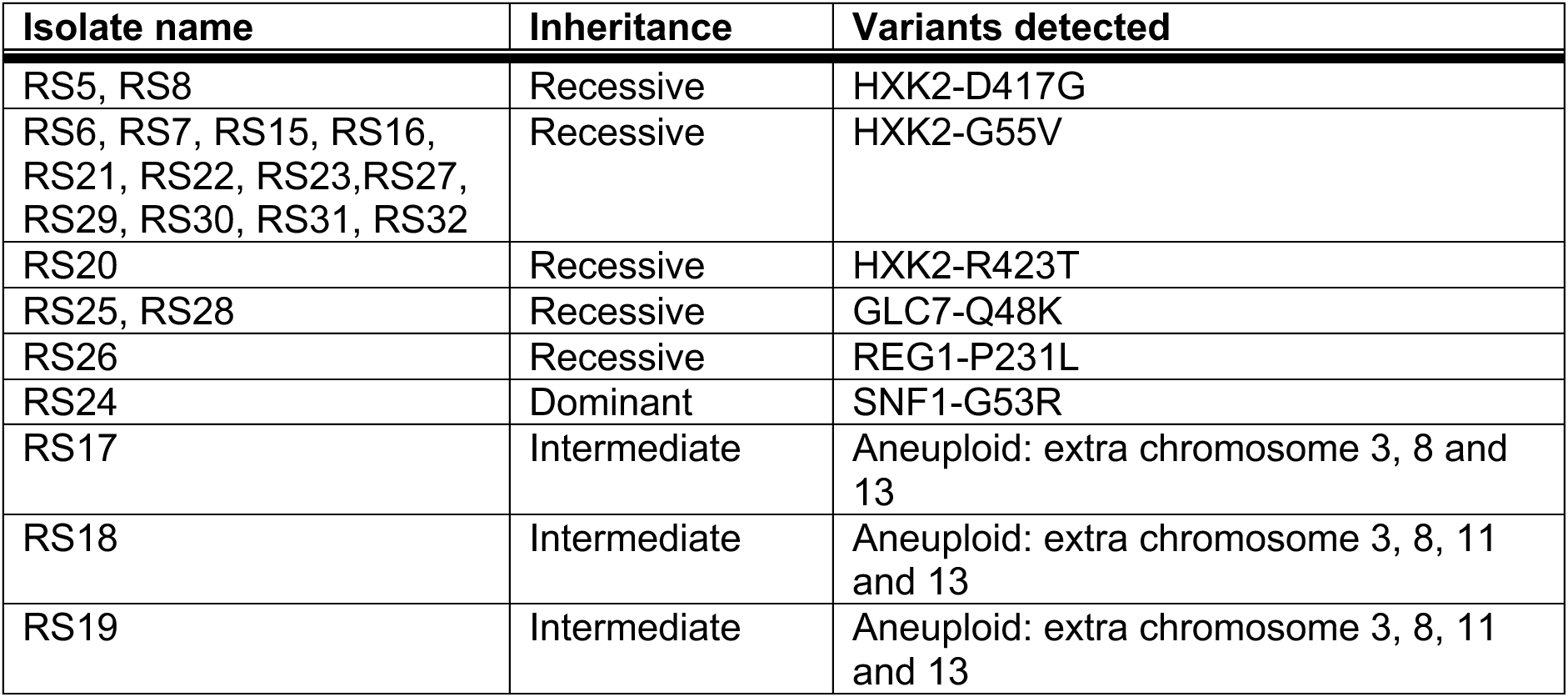
Genomic Sequencing Results

In order to confirm that these candidate mutations did in fact cause 2DG resistance, we engineered plasmids to contain these mutations and introduced these plasmids into strains lacking the cognate gene. In all cases, the candidate mutations when present on low-copy plasmids produced the same 2DG resistance as was observed when the mutations were in the chromosomes (not shown). After confirming that these single mutations were sufficient to confer the 2DG-resistant phenotypes, we sought to characterize each mutation further and uncover how they play a role in in 2DG resistance.

### Recessive, missense alleles in *HXK2* gene inhibit catalytic activity and confer 2DG resistance

We used oligonucleotide-directed mutagenesis to generate several versions of *HXK2* all with a C-terminal epitope tag. We made the three missense alleles isolated in this screen (*hxk2*-G55V, *hxk2*-D417G, and *hxk2*-R423T). We also tagged two additional alleles of *HXK2* that had been previously reported to separate the catalytic and gene regulatory functions of Hxk2 (15). These alleles are referred to as *HXK2*-wca (without catalytic activity) and *HXK2*-wrf (without regulatory function). We previously reported that the *HXK2-wca* mutation conferred 2DG-resistance, while the *HXK2-wrf* mutation did not confer 2DG resistance (5). Finally, we created the *hxk2*-D211A allele, since the aspartate at 211 is a catalytic residue and the substitution of this residue with alanine is known to severely compromise catalytic activity (16).

The three *HXK2* alleles isolated in this screen were found to be highly 2DG resistant (Fig 1A). Complete deletion of the *HXK2* gene conferred the greatest resistance to 2DG with the *hxk2*-G55V and *hxk2*-D211A alleles following close behind. Consistent with our earlier studies (5), the *HXK2-wca* allele conferred 2DG resistance while the *HXK2-*wrf allele did not. To assess Hxk2 protein expression and catalytic activity, we utilized a strain lacking all three hexokinase genes (*hxk1Δ hxk2Δ glk1*Δ). This strain can grow on galactose but is unable to grow by fermentation of glucose (Fig. S2). The triple hexokinase delete strain was transformed with low-copy plasmids expressing epitope-tagged wild type or one of the six alleles of *HXK2* described above. A western blot of protein extracts from cells grown on galactose demonstrates that all proteins were expressed as full-length proteins although the absolute level of Hxk2 protein accumulation showed some variation (Fig 1B).

**Fig 1.**
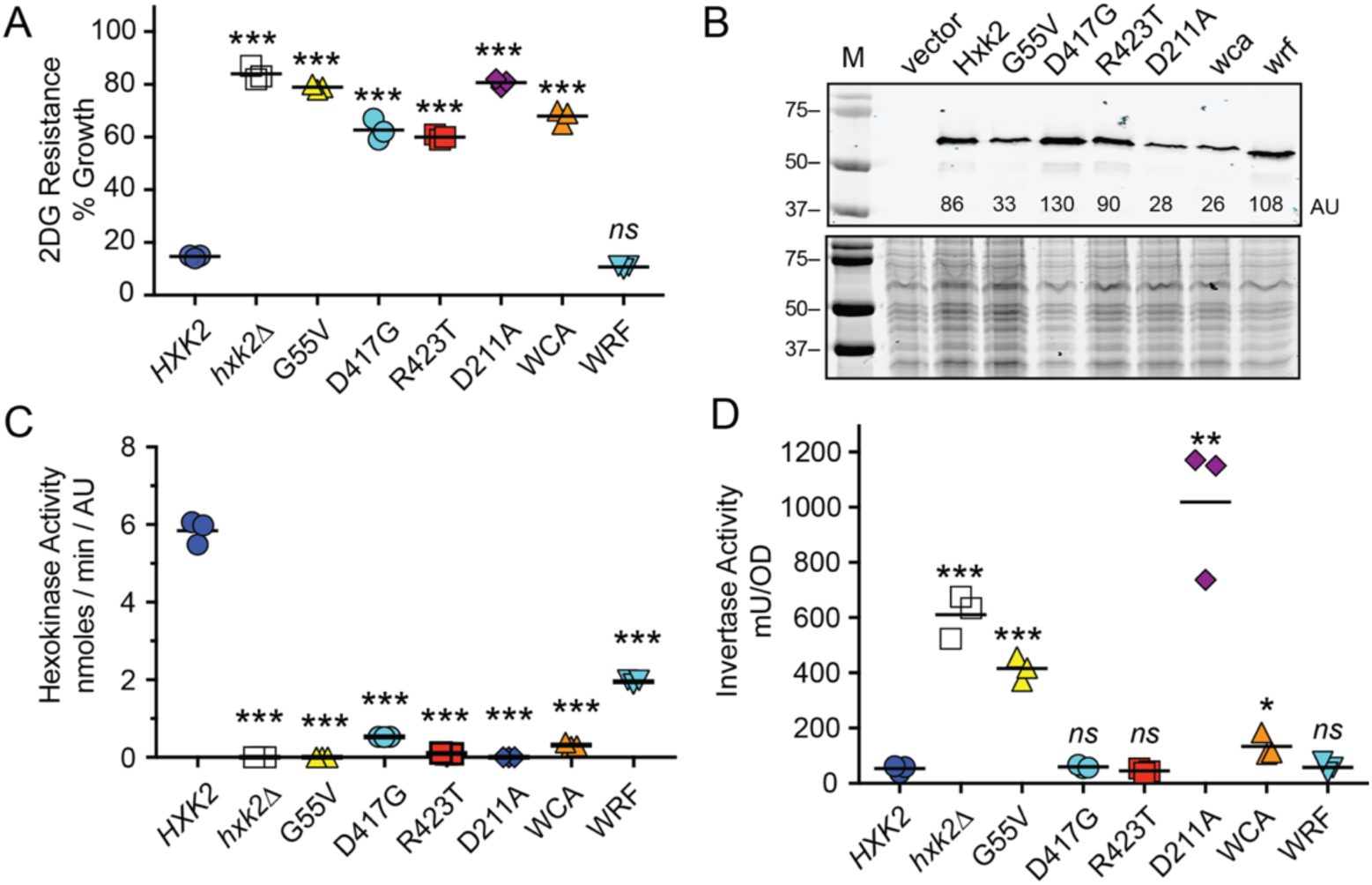
Amino acid substitutions in Hxk2 that confer resistance to 2-deoxyglucose. **A.** 2-Deoxyglucose resistance assay in *hxk2*Δ cells transformed with low-copy plasmids encoding wild type (WT) *HXK2, HXK2* with the indicated amino acid substitutions, the wca and wrf alleles (15) or empty vector (*hxk2*Δ). Three independent cultures were grown in media with and without 0.1% 2DG with percent growth plotted relative to growth in the absence of 2DG. Individual data points are shown with the mean indicated with a solid bar. Values statistically different from wild type are indicated. **B.** Western blot of Hxk2 proteins tagged with three copies of the V5 epitope at the C-terminus. Quantitation of the western signal in arbitrary units (AU) is shown. As a control for equal loading, Coomassie stained gel of the same extracts is shown below. **C.** Catalytic activity of hexokinase was measured in protein extracts of cells lacking all three hexokinase genes (*hxk1Δ hxk2Δ glk1*Δ). Cells were transformed with the same low-copy plasmids in A. Activity is expressed in nmoles per minute normalized to the amount of protein in the extract as determined by western blotting. Values statistically different from wild type are indicated. **D.** Invertase activity measured in three independent transformants of *hxk2*Δ cells with the same plasmids used in A. Values statistically different from wild type are indicated.

Hexokinase II is reported to have two distinct activities, a catalytic activity and a gene regulatory activity. We measured both of these activities in the Hxk2 mutants under study. First, we used a coupled *in vitro* enzyme assay system (17, 18) to measure the catalytic activity of each Hxk2 enzyme (Fig 1C). Catalytic activity was normalized to the amount of Hxk2 protein as quantified by western blotting (Fig 1B) using the arbitrary units (au) shown. Extracts from cells lacking any hexokinase and from cells expressing the D211A allele did not contain any detectable hexokinase activity. All the mutant Hxk2 enzymes isolated in this study showed severely compromised catalytic activity ranging from 3% to 0.3% of that observed in cells expressing wild type Hxk2. The catalytic activity and the 2DG resistance phenotype show a very strong inverse correlation. The mutant enzyme with the most catalytic activity (hxk2-wrf) confers the least 2DG resistance, while the mutants with the least catalytic activity (hxk2-G55V and hxk2-D211A) showed the highest levels of 2DG resistance. One curious observation about the three yeast hexokinase isozymes (*HXK1*, *HXK2* and *GLK1*) is that only deletion of *HXK2* confers 2DG resistance (Fig S2). This observation could be explained if the Hxk2 enzyme was the only one of these isozymes capable of phosphorylating 2DG. We tested this in our *in vitro* assay and found that both Hxk1 and Hxk2 were able to phosphorylate 2DG *in vitro*. In fact, the Hxk1 enzyme had a higher affinity (lower Km) and higher Vmax for 2DG than the Hxk2 enzyme (Table 2 and Figs S4 and S5). The Glk1 enzyme exhibited a much higher affinity (lower Km) for glucose than either Hxk1 or Hxk2 but was not able to detectably phosphorylate 2DG in our *in vitro* assay (Fig S6). We suspect that Glk1 is able to phosphorylate 2DG to some extent *in vivo* since introduction of *GLK1* to the triple hexokinase mutant confers sensitivity to 2DG when growing on glycerol/ethanol media (Fig S2B).

**Table 2.** Kinetic analysis of hexokinases.

| Hexokinase | Km<br>Glucose<br>(mM) | Vmax<br>Glucose<br>(nmole/min) | Km<br>2DG<br>(mM) | Vmax<br>2DG<br>(nmole/min) |
| --- | --- | --- | --- | --- |
| *HXK1* | 0.11 | 0.42 | 0.12 | 0.26 |
| *HXK2* | 0.19 | 0.26 | 0.28 | 0.075 |
| *GLK1* | 0.055 | 0.17 | ND<sup>a</sup> | ND<sup>a</sup> |

<sup>a</sup>ND, not detected

In order to assess the gene regulatory function of Hxk2, we measured invertase expression in cells grown on high glucose. Previous studies have shown that invertase expression is repressed in wild-type cells growing on high glucose but is significantly derepressed (induced) when the *HXK2* gene has been deleted or mutated (2, 19). We found that the cells containing the lowest levels of Hxk2 catalytic activity (*hxk2*Δ, *hxk2-G55V* and *hxk2-D211A*) had the greatest derepression of invertase (Fig 1D). Surprisingly, some *HXK2* alleles (*hxk2-D417G*, *hxk2-R423T* and *hxk2-wca*) showed normal levels of invertase repression even though they were all severely compromised for hexokinase catalytic activity (Fig 1B). The Hxk2-wrf enzyme has been reported to lack gene regulatory activity (15), as judged by the derepression of invertase. We have been unable to reproduce that finding. Cells with the *hxk2-wrf* allele showed wild-type levels of invertase repression in our assay (Fig 1D). We have sequenced the *hxk2-wrf* and confirmed that it contains the deletion of residues 6-15 that are reported to cause a loss of regulatory function. Our results are consistent with those reported from the Botstein lab showing that deletion of amino acids 1-15 in Hxk2 had little effect on invertase expression (17). Our results reported here and those reported earlier (18) indicate that *HXK2* mutations do not have a direct correlation between catalytic activity and catabolite repression. In contrast and consistent with our earlier study (5), we find that loss of catalytic activity of Hxk2 strongly correlates with 2DG resistance (Table 3).

**Table 3.**
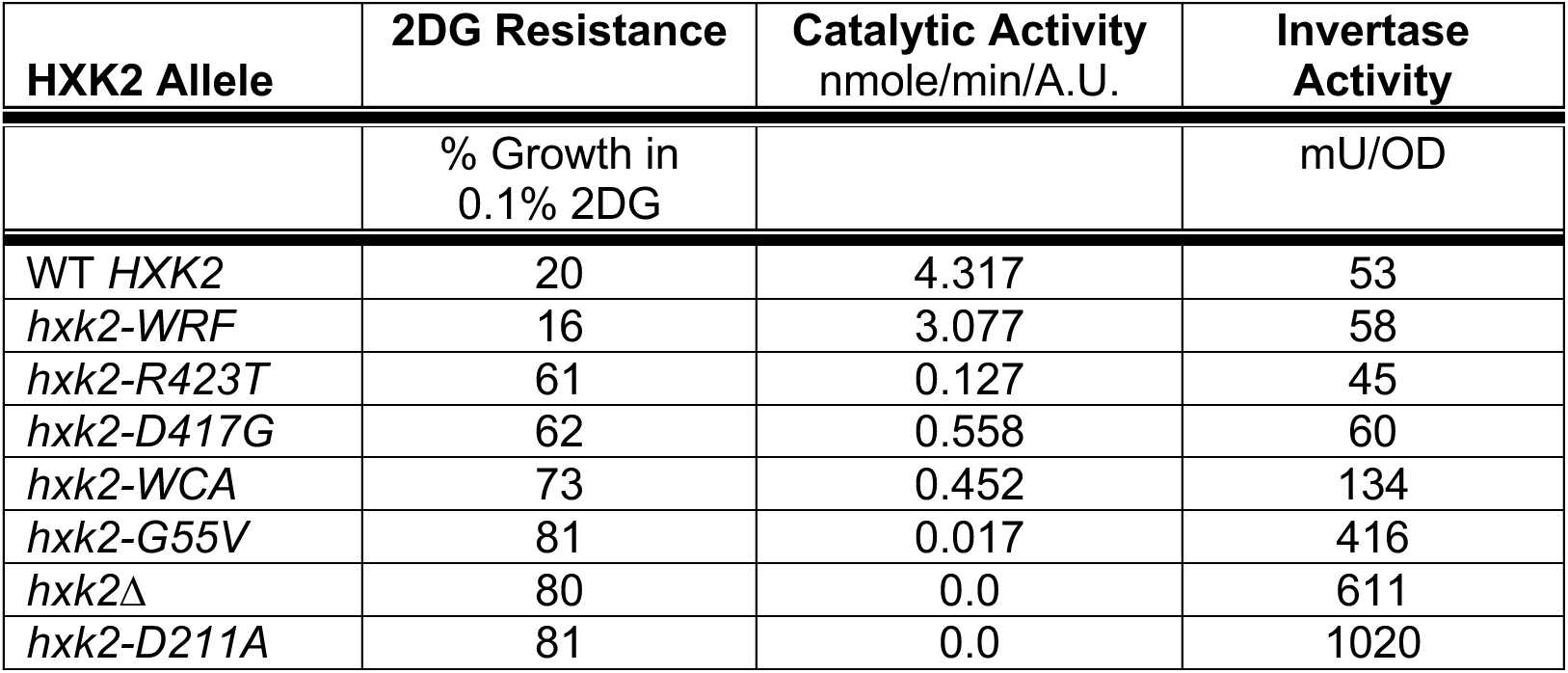
Activity of *HXK2* and variants.

### Recessive, missense alleles in PP1 phosphatase genes *REG1* and *GLC7* confer 2DG resistance

Yeast PP1 phosphatase is composed of the catalytic subunit Glc7 associated with a number of alternative regulatory subunits (11). The Glc7/Reg1 complex is the form of PP1 that participates in glucose repression by dephosphorylating the kinase Snf1 (20–22) and the transcription factor Mig1 (21, 23). The *reg1-P231L* mutation found in our screen confers significant 2DG resistance in yeast when reconstituted on a low-copy CEN plasmid in a *reg1*Δ strain (Fig 2A). For comparison, we also measured 2DG resistance in cells lacking the *REG1* gene (*reg1*Δ) and in cells expressing a variant of Reg1 (*reg1-IF*) that cannot bind to Glc7 (24, 25). Cells lacking Reg1 or expressing Reg1-IF are almost completely resistant to 2DG. Cells lacking Reg1 are defective in catabolite repression and express high levels of invertase even in the presence of glucose (Fig 2B). The Reg1*-*P231L protein did not show any defect in invertase repression and was present in cells at levels comparable to wild type Reg1 as judged by western blotting (Fig 2C). The Reg1 protein binds to the Snf1 kinase complex as judged by immunoprecipitation and two-hybrid analysis (25, 26). While we did notice some impairment in the interaction of Snf1 with the Reg1-P231L mutant by the yeast two-hybrid assay (Fig S3), no significant effect on Snf1 dephosphorylation was detected by western blotting with antibodies specific for the phosphorylated form of Snf1 protein (Fig 2D).

**Fig. 2.**
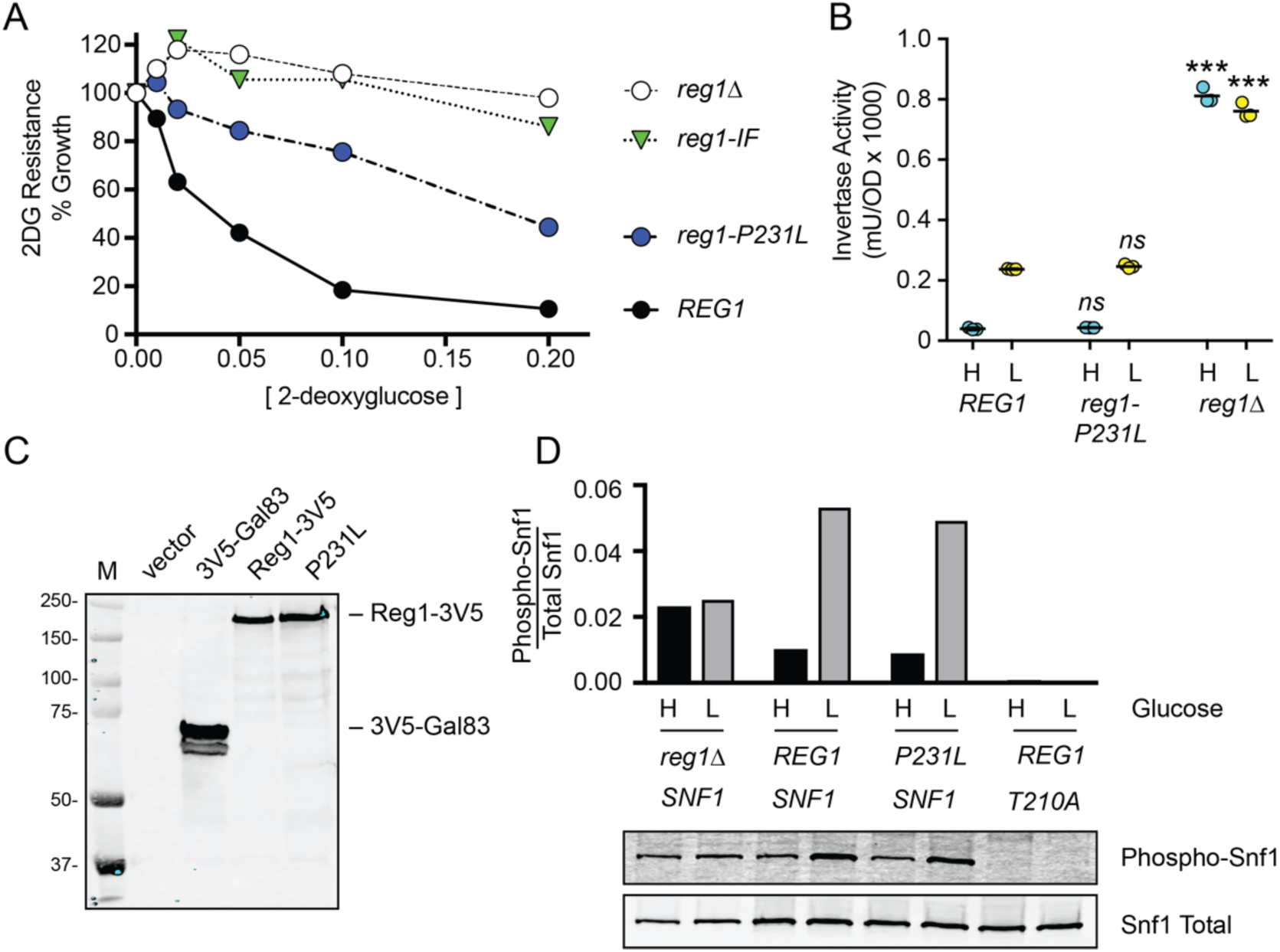
Amino acid substitution, Reg1-P231L, confers 2DG resistance. **A.** 2DG resistance assay of *reg1*Δ cells transformed with empty vector or plasmids expressing wildtype Reg1, Reg1-P231L or the Reg1-IF mutation known to disrupt interaction with Glc7 (24). **B.** Invertase assays using cells from three independent cultures grown in high glucose (H) or two hours after shifting to low glucose (L) as indicated. The *REG1* genotype shown below. Individual data points are shown with the mean indicated with a solid bar. Values of the mutants were compared to wild type and statistical significance is shown. **C.** Western blot using V5 epitope-tagged proteins expressed from low-copy number plasmids. **D.** Western blot of HA-tagged Snf1 protein kinase using antibodies that detect total Snf1 protein or Snf1 protein phosphorylated on threonine 210. Quantitation of the western signals is shown in the bar graphs above the blot images.

The *GLC7* gene encodes the catalytic subunit of the PP1 phosphatase and is essential for viability (27). To confirm that the *GLC7-Q48K* allele conferred 2DG resistance, we transformed a diploid heterozygous strain (*GLC7/glc7*Δ) with a low-copy plasmid encoding an HA-tagged *GLC7* with and without the Q48K mutation. After sporulation, we recovered viable haploid cells with both plasmid encoded *GLC7* genes and the chromosomal *glc7Δ::KAN* allele. We assayed both the wild type and the Q48K mutant strains and confirmed that the *glc7-Q48K* allele conferred 2DG resistance (Fig 3A). This mutation did weaken the yeast two-hybrid interaction with Reg1 (Fig S3), but had no discernible effect on the phosphorylation of Snf1 (Fig 3B). The *glc7-Q48K* mutation had a large effect on the regulation of invertase activity (Fig 3C). In cells expressing wild type Glc7-3HA, invertase expression was repressed in high glucose and induced when cells are shifted to low glucose. In contrast, cells expressing Glc7-3HA with the Q48K mutation showed high levels of invertase expression in both high and low glucose. Repression of invertase expression in high glucose requires dephosphorylation of the Mig1 transcriptional repressor (23). The phosphorylation state of Mig1 is readily discernable, since phosphorylation of Mig1 causes significant reduction in gel mobility that was reversed by phosphatase treatment (28). We tested whether any of the 2DG-resistant mutations in the PP1 phosphatase affected the phosphorylation state of Mig1. In wild-type cells, the phosphorylation of Mig1 and its associated reduction in gel-mobility was readily detected upon shifting cells to low glucose (Fig 3D). A similar pattern of Mig1 phosphorylation was observed for cells containing the *glc7-Q48K* and *reg1-P231L* alleles. Thus, the two missense mutations in the PP1 complex had no detectable effect on Mig1 phosphorylation and yet caused very different effects on catabolite repression.

**Fig 3.**
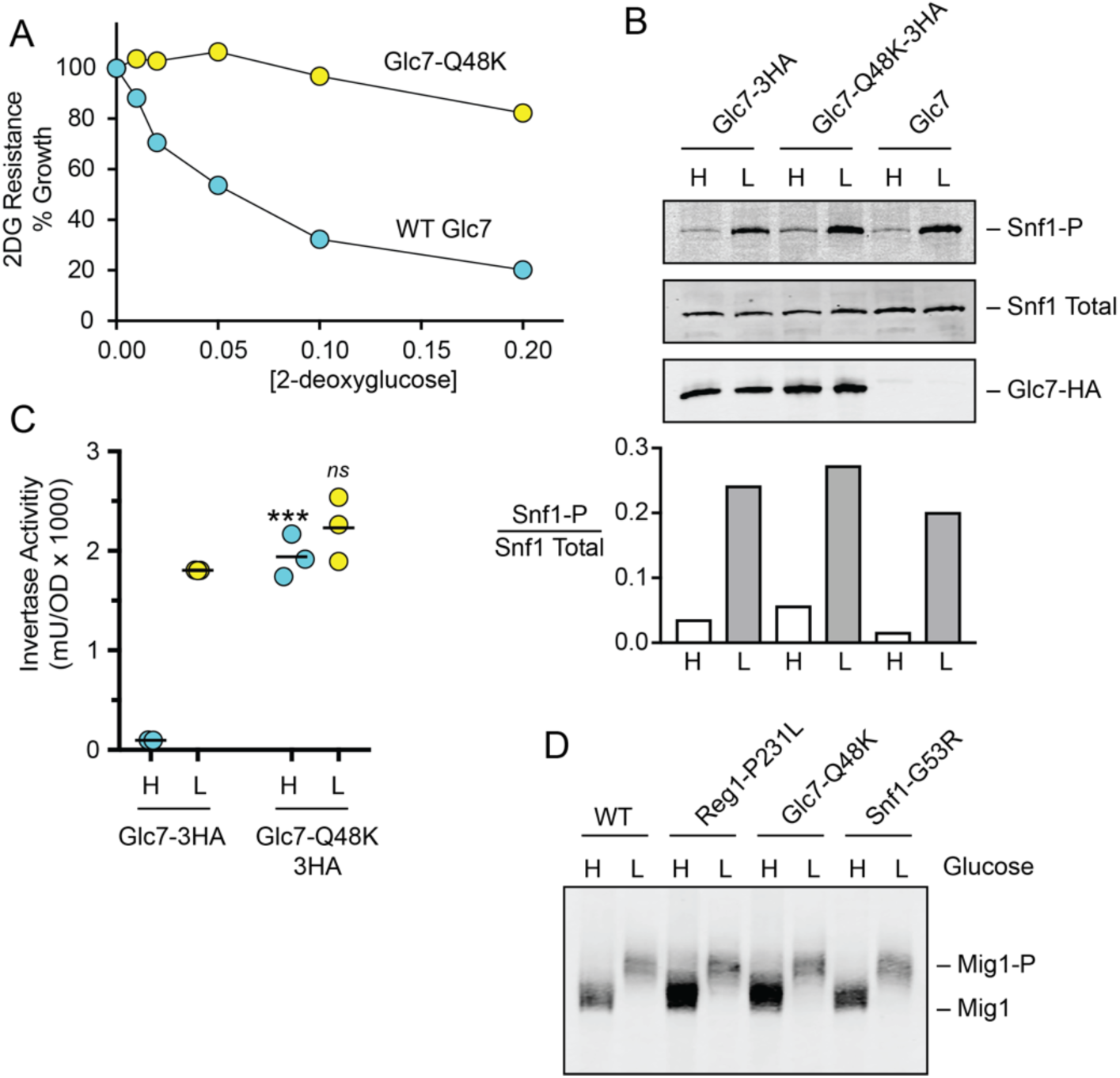
Amino acid substitution, Glc7-Q48K, confers 2DG resistance. **A.** 2DG resistance assay in *glc7*Δ cells containing plasmids to express HA-tagged wildtype Glc7 or HA-tagged Glc7-Q48K. **B.** Western blot of Snf1 kinase using antibodies to detect total Snf1 protein, Snf1 protein phosphorylated on threonine 210, or HA-tagged Glc7 protein. A plot of the ratio of phospho-Snf1 to total Snf1 is shown below. **C.** Invertase activity of three independent cultures grown in high glucose (H) or two hours after shifting to low glucose (L). Individual data points are shown with the mean indicated with a solid bar. Values statistically different from wild-type cells in the same glucose condition are indicated. **D.** Western blot of Mig1-3HA protein in cells grown in high (H) glucose or two hours after shifting to low (L) glucose. Relevant genotypes are shown above. Migration of Mig1 and phosphorylated Mig1 are indicated.

### Dominant allele of *SNF1* confers 2DG resistance

In our screen for 2DG-resistant mutations, we characterized 28 mutations. Twenty-seven were recessive mutations and one was dominant. When we sequenced the genome containing the one dominant allele, we identified the missense allele, *SNF1-G53R,* as the primary candidate responsible for 2DG resistance (Table 2). This allele of *SNF1* has been isolated previously as a dominant mutation that improves Snf1 kinase function in the absence of its gamma subunit, Snf4 (14). To confirm that the *SNF1-G53R* mutation conferred 2DG resistance, *snf1*Δ cells were transformed with plasmids expressing wild type *SNF1* with a triple HA tag at the C-terminus (20), the *SNF1-G53R* allele, or empty vector. As we have previously shown, cells lacking a *SNF1* gene (*snf1*Δ) are hypersensitive to 2DG (5). Cells with the *SNF1-G53R* were resistant to 2DG relative to wild-type cells (Fig 4A), consistent with the idea that *SNF1-G53R* is a hyperactive *SNF1* allele. The phosphorylation status of the Snf1 kinase activation loop is a good proxy for determining Snf1 kinase activity (20). We next examined whether the G53R substitution affected the phosphorylation status of the Snf1 activation loop using antibodies specific for Snf1 phosphorylated on threonine 210. The Snf1 kinase with the G53R substitution showed normal basal levels of threonine 210 phosphorylation in cultures grown in high glucose and somewhat higher levels after cultures were shifted to low glucose (Fig 4B). However, when we analyzed the effect of this mutation on gene expression, we found that invertase expression was not significantly affected by this mutation (Fig 4C). A similar result with the *SNF1-G53R* allele was reported previously (14). A known substrate of the Snf1 kinase is the transcriptional repressor Mig1. Cells expressing Snf1-G53R did not show altered phosphorylation of the Mig1 protein (Fig 3D). Thus, the mechanism by which Snf1-G53R confers 2DG resistance is not likely to involve changes in Mig1-regulated gene expression.

**Fig 4.**
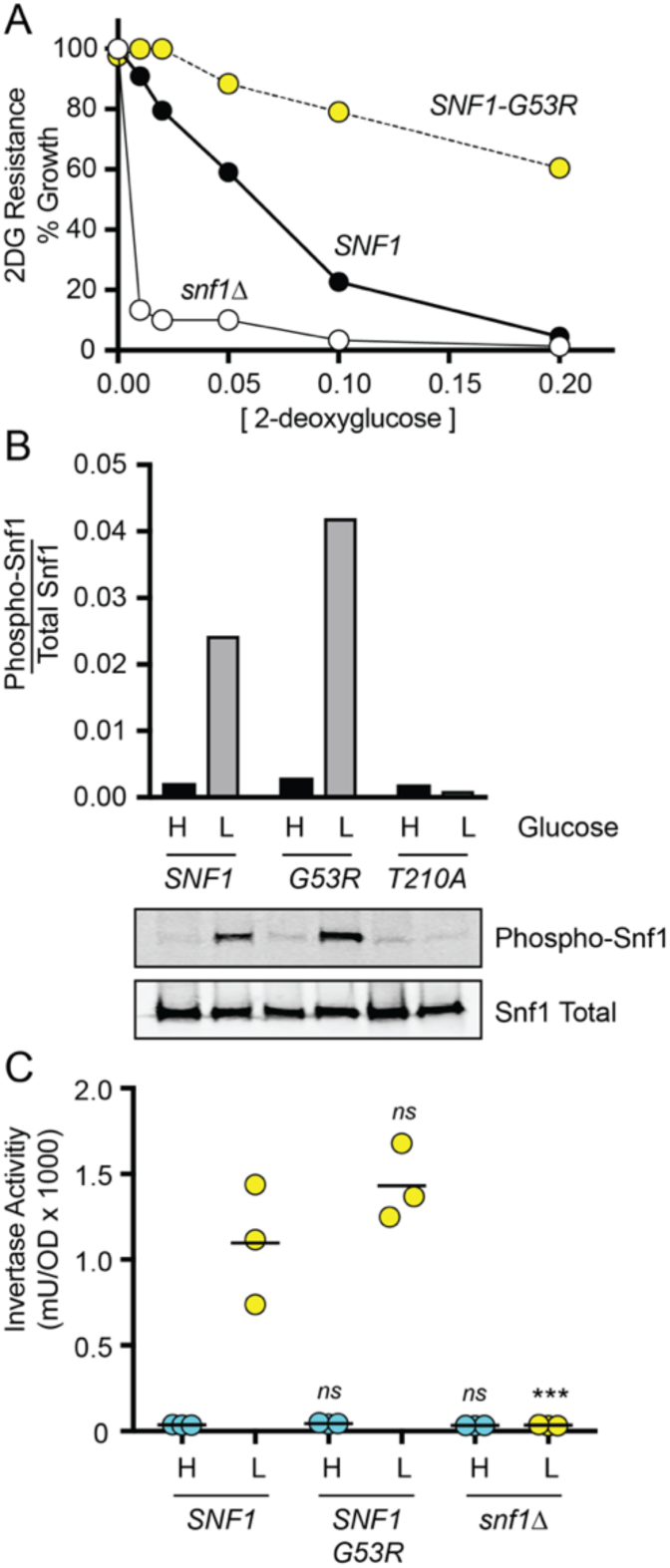
Amino acid substitution, Snf1-G53R, confers 2DG resistance. **A.** 2DG resistance assay in *snf1*Δ cells transformed with plasmids expressing either HA-tagged wild-type Snf1, HA-tagged Snf1-G53R, or no Snf1 (*snf1*Δ). **B.** Western blot of HA-tagged Snf1 protein kinase using antibodies that detect total Snf1 protein or Snf1 protein phosphorylated on threonine 210. Quantitation of the western signals is shown in the bar graphs above the blot images. **C.** Invertase activity was measured in three independent cultures grown in high glucose (H) or two hours after shifting to low glucose (L). Values statistically different from wild-type values in the same glucose condition are indicated.

### Mutations in Hxk2 and Snf1 pathway genes protect Hxt3 from endocytosis

In our earlier study of the mechanism of 2DG toxicity, we found that 2DG promoted the endocytosis of the high-capacity glucose transporters Hxt1 and Hxt3 (7). The endocytosis of these glucose transporters was dependent on the arrestins Rod1 and Rog3 and was regulated by the Snf1 kinase. The wild-type, 2DG-sensitive yeast strain (MSY1333; Table 4) that we used to select new spontaneous mutations contained an integrated copy of *HXT3-GFP* gene allowing us to track the localization of this glucose transporter by fluorescence microscopy. We measured the localization of Hxt3-GFP in our wild-type strain either grown in glucose or 2 hours after exposure to 2DG. Consistent with our earlier report, we found that 2DG promoted endocytosis of Hxt3-GFP (Fig 5A). This effect was quantified by measuring Hxt3-GFP fluorescence in individual cells and plotting the ratio of the fluorescence intensity at the plasma membrane relative to that of the vacuole (Fig 5B). When we analyzed Hxt3-GFP fluorescence in cells containing 2DG-resistant mutations, we found a significant increase in the plasma membrane localization of Hxt3-GFP for the cells expressing Hxk2-D417G, Hxk2-R423T, Snf1-G53R, Glc7-Q48K and Reg1-P231L. Curiously, the cells expressing Hxk2-G55V did not show any significant retention of Hxt3-GFP at the plasma membrane. Thus, the three Hxk2 mutations isolated in this study appear to have different effects in HXT3-GFP localization. The finding that all three mutations in the Snf1 pathway (*SNF1-G53R, glc7-Q48K and reg1-P231L)* promoted retention of Hxt3-GFP at the plasma membrane is consistent with our earlier finding that the Snf1 kinase plays an important role in regulating endocytosis of the Hxt3 protein. The mechanism by which some of the *HXK2* alleles regulated HXT3-GFP localization is not yet known.

**Table 4.**
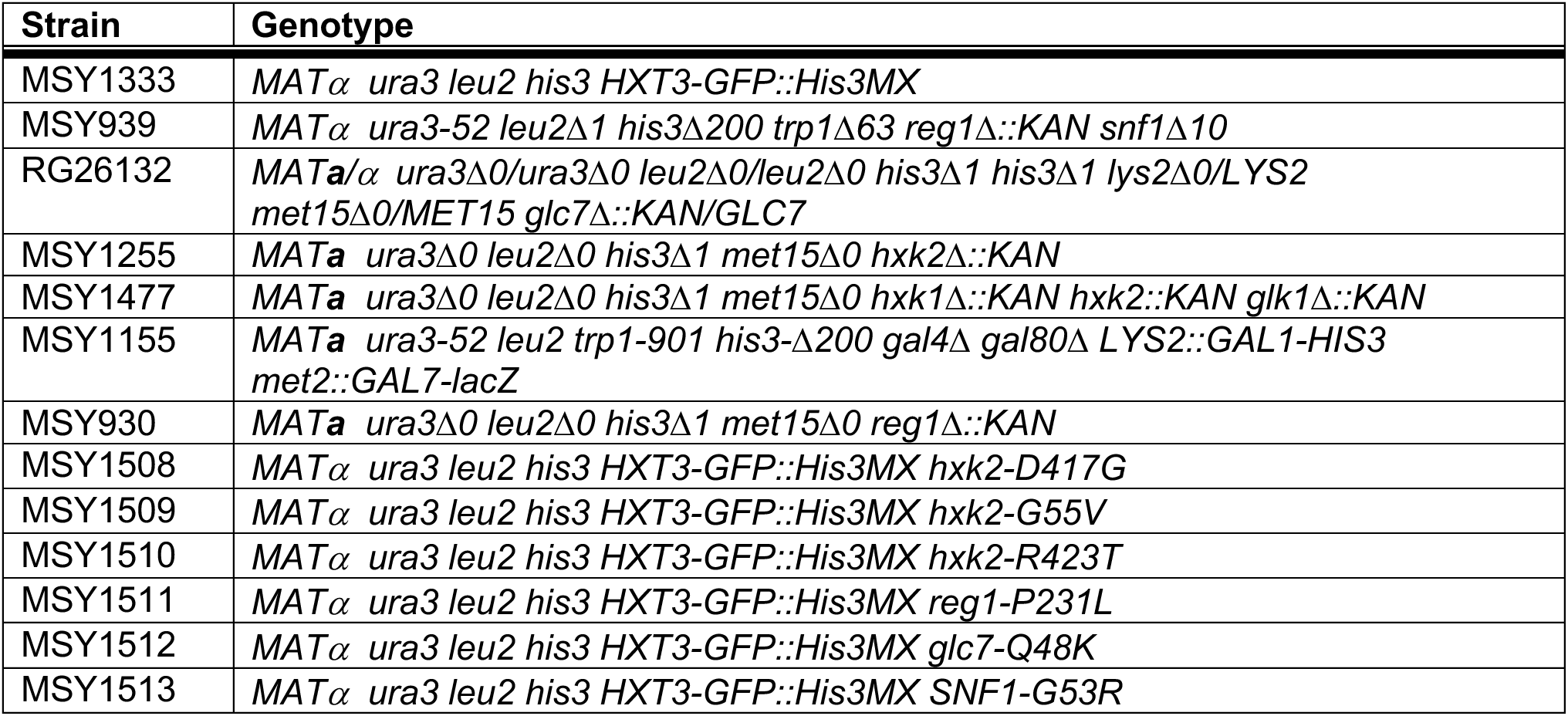
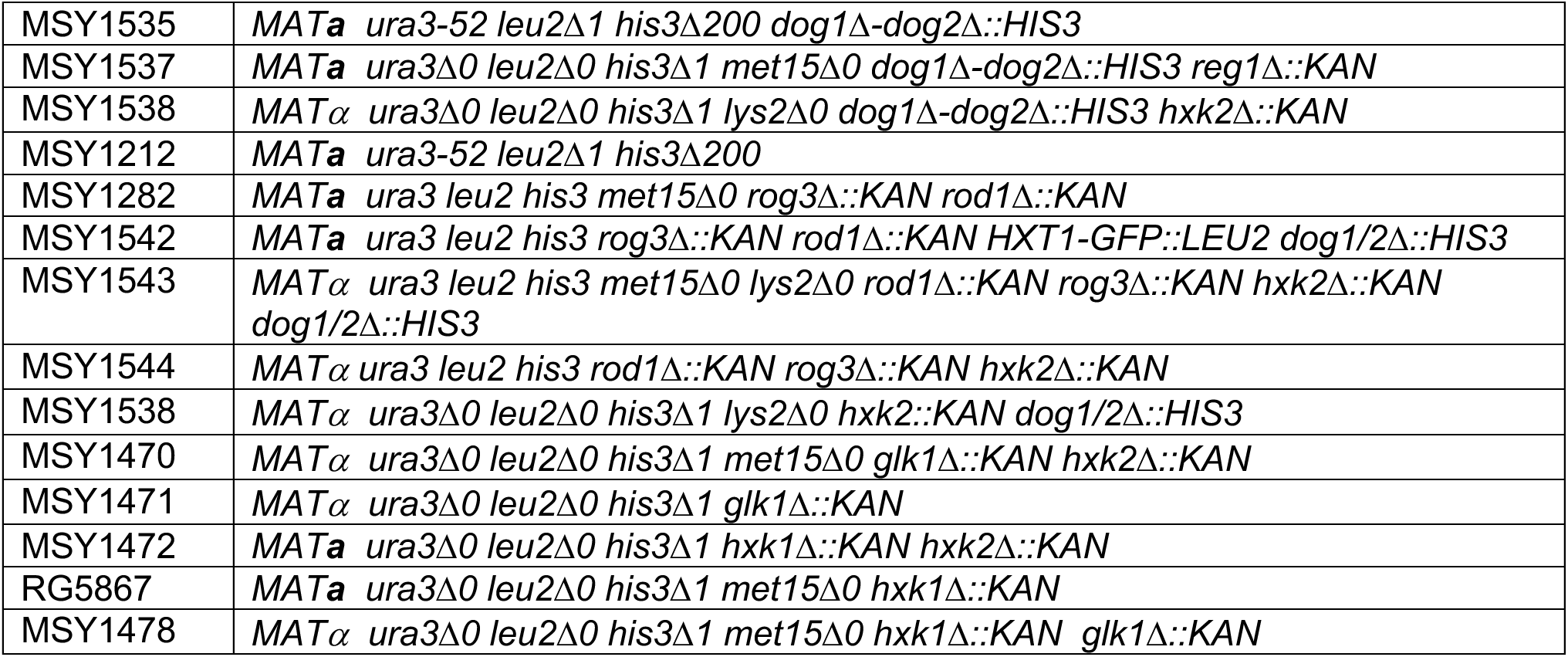
Yeast strains

**Fig. 5.**
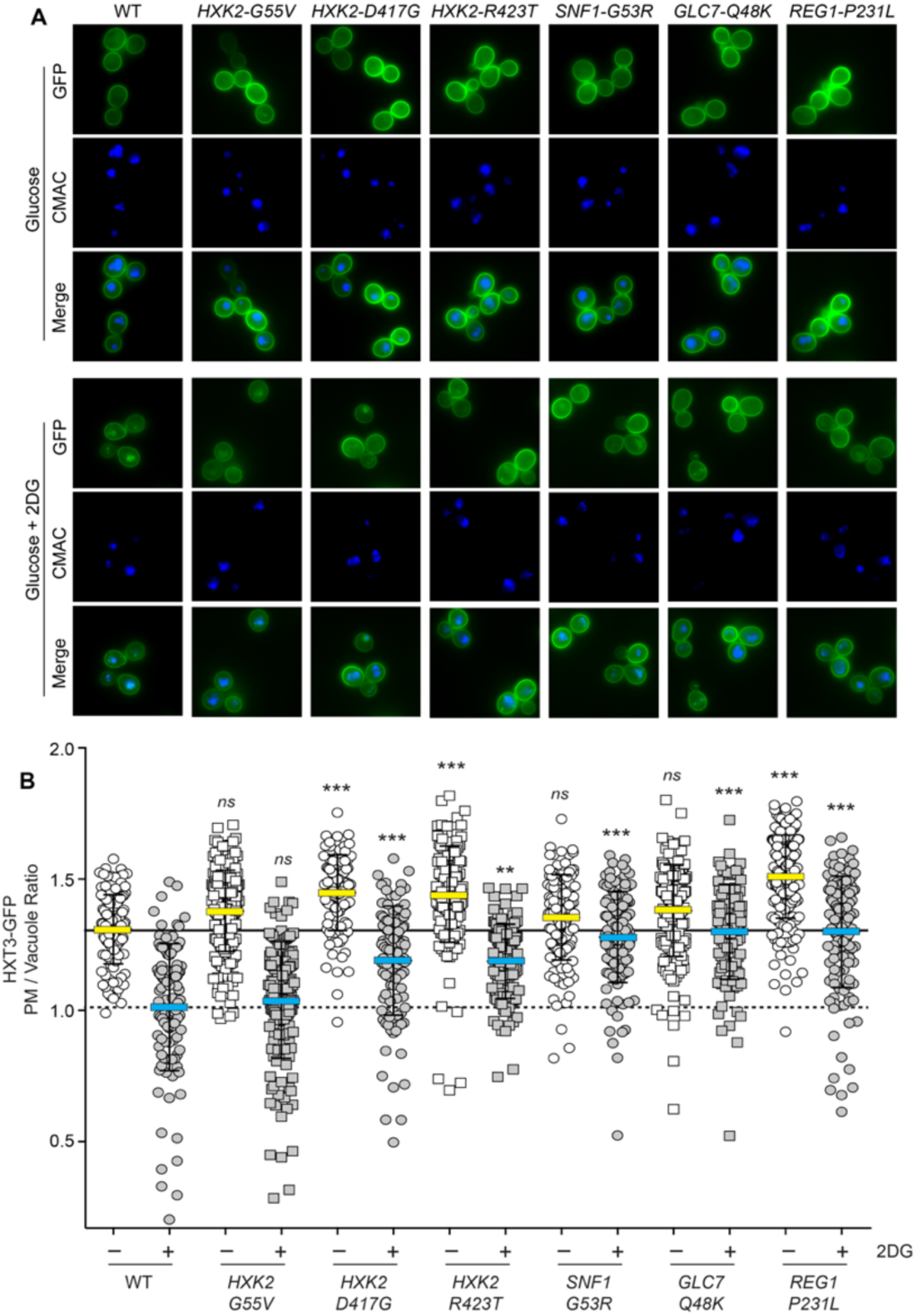
HXT3-GFP localization. **A.** Wild-type cells (WT) or cells with the indicated mutations with integrated Hxt3-GFP were stained with CMAC blue to identify the vacuole. Images were captured during growth on glucose or two hours after addition of 2DG. **B.** Quantitation of GFP fluorescence for at least 150 cells is plotted for each cell as the ratio of plasma membrane (PM) fluorescence to vacuolar fluorescence. The mean value for cells grown in glucose is marked with a yellow bar with the mean value for wild-type cells indicated as a solid reference line across the graph. The mean value for cells treated with 2DG is marked with a blue bar with the mean value for wild-type cells treated with 2DG is indicated with a hashed line as a reference value across the graph. Fluorescence values statistically different from wild type, where all untreated cells are compared to untreated WT and all 2DG-treated cells are compared to the 2DG-treated WT control.

### Transcriptional response to acute carbon source stress

In order to better understand the cell’s transcriptional response to carbon source stress, we measured the abundance of mRNAs in cells using RNAseq. Total RNA was harvested from wild-type cells growing in synthetic complete media with 2% glucose as the carbon source or from cells shifted for two hours to either low glucose (0.05% glucose) or 2% glucose plus 0.1% 2DG. RNA samples were prepared from at least four independent cultures and at least fifty million reads were mapped for each sample. Abundance of each mRNA was determined using kallisto software (29) and is expressed as transcripts per million mapped reads (tpm). We plotted the mean expression of the mRNA under both conditions on the x-axis and on the y-axis, the log2 ratio of the mean tpm values for the two conditions (Fig 6A). This allows for simultaneous visualization of the change in expression and the relative abundance of an mRNA. One immediate transcriptional response of cells undergoing carbon source stress was the reduction in the abundance of ribosomal protein mRNAs (red circles). Both low glucose stress and 2DG stress cause an average reduction in ribosomal mRNAs by about 10-fold (Fig 6B). We specifically examined the *HXT* genes, since their abundance and localization has been linked to 2DG resistance (7). When cells were shifted to low glucose, the mRNA abundance for the high capacity transporters (*HXT1, HXT3 and HXT4*) decreased dramatically, while the high affinity transporters (*HXT5* and *HXT6*) showed increased mRNA abundance (yellow circles). This response to limited glucose abundance has been reported previously using lacZ-reporter assays (30, 31). In contrast, when cells are exposed to 2DG, they responded by decreasing the abundance of the highly expressed *HXT* genes but do not induce high level expression of any of the high affinity *HXT* genes. Therefore, cells exposed to 2DG respond both by decreasing the expression of *HXT* mRNA and by removing *HXT* proteins from the plasma membrane (7). The deoxyglucose phosphatase genes (*DOG1* and *DOG2*) have also been implicated in the response to 2DG (10). We found that the *DOG* transcriptional response to these two stressors was quite distinct (Fig 6A and 6B; white diamonds) and is discussed in more detail below. Examining the transcriptional response globally shows considerable overlap between these two carbon source stressors (Fig 6C and 6D). We arbitrarily defined a significant transcriptional response as a 4-fold change in mRNA abundance (log2 ratio greater than 2 or less than −2) with statistical significance (p <0.01). With these cut-offs, we found that over 450 genes showed a significant transcriptional response and were co-regulated under these stress conditions. Based on gene ontology, co-regulated genes were enriched for processes that might be expected in response to carbon source stress, including increased abundance in genes associated with mitochondrial function, carbohydrate metabolism and cell wall metabolism.

**Fig 6.**
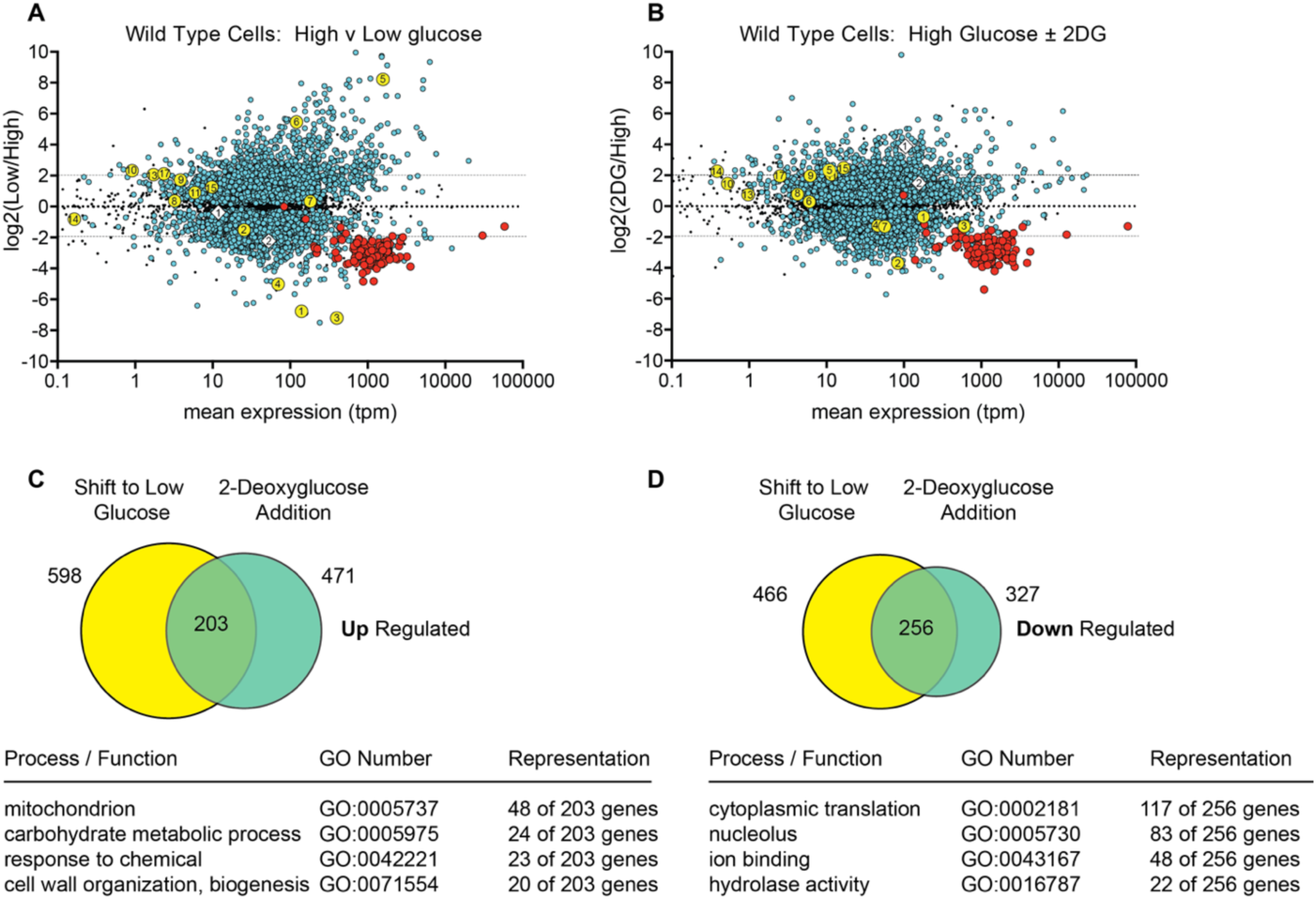
RNAseq analysis of wild-type cells undergoing carbon source stress. RNA was extracted from four independent yeast cultures of wild-type cells (MSY1212) grown to mid-log phase in synthetic complete media with 2% glucose and either two hours after shifting to 0.05% glucose **(A)** or two hours after addition of 2-deoxyglucose to 0.1%. **(B)**. RNA abundance was quantified and is displayed here for 5917 mRNAs as the log2 ratio of the mean expression values (tpm) on the y-axis and the mean expression values under both conditions on the x-axis. Expression levels that show a statistically significant change (p < 0.01) are shown as colored dots. Those not meeting this threshold are shown as smaller black dots. Ribosomal protein genes are shown as red circles. Hexose transporters are plotted as yellow circles with the number of the *HXT* genes shown. The *DOG1* and *DOG2* mRNAs are shown as white diamonds. Genes up-regulated by 4-fold **(C)** or down-regulated by 4-fold **(D)** and meeting the statistical significance threshold for each dataset are plotted in a Venn diagram. Significant gene ontology terms (GO) for the genes in the intersecting sets are shown.

### Competitive fitness of strains with 2DG-resistant alleles

At the outset of this selection for 2DG resistance, we expected to recover *REG1* and *HXK2* loss of function alleles, since the deletion of either of these genes is known to confer 2DG resistance (5). However, we did not recover any nonsense alleles in our screen (Table 2). This result is understandable for *GLC7*, since this is an essential gene, and nonsense alleles would not be viable, and could be anticipated for *REG1* since *reg1*Δ cells grow very poorly are at a selective disadvantage overall (32). However, cells lacking *HXK2* grow normally on glucose and exhibit resistance to 2DG (Fig S2A), and therefore nonsense mutations in *HXK2* were expected but not recovered. We noticed that the *REG1-P231L* allele did not confer a slow growth phenotype. We examined the relative fitness of all the 2DG resistance alleles recovered in this screen by growing strains in competitive cultures in synthetic complete media with glucose or glucose plus 0.1% 2DG. After competitive growth, the representation of each genotype was determined based on different auxotrophic markers. The relative fitness index (33) of each pair of strains was plotted on a log2 scale (Fig S7). For the *HXK2* alleles, we found that complete deletion of *HXK2* exacted a very small fitness cost compared to wild type for cells growing on glucose and conferred a large fitness advantage to cells growing in the presence of 2DG. The missense alleles of *HXK2* all showed a similar advantage in media with 2DG with very low cost on glucose. A very different fitness landscape is seen with *REG1* alleles. Deletion of *REG1* exacts a large fitness cost for cells growing on glucose (over 4-fold). Cells with *reg1*Δ or the *reg1-P231L* allele showed a large increase in fitness when challenged with 2DG compared to wild type *REG1* cells. Interestingly, the *reg1-P231L* allele showed increased fitness compared to the *reg1*Δ allele in both glucose and glucose plus 2DG media. The competitive advantage of the *reg1-P231L* allele over the *reg1*Δ allele explains why the *reg1-P231L* allele was isolated in preference to loss of function alleles that come with a strong fitness disadvantage. When we examined the relative fitness of the *SNF1* alleles, we found that the loss of *SNF1* came with a reduced fitness under both media conditions. The *SNF1-G53R* allele confers increased fitness compared to wild type when challenged with 2DG with almost no fitness cost to cells growing on glucose. The *SNF1-G53R* conferred an advantage to cells compared to the *snf1*Δ on both media. Finally, we compared the *GLC7* wild-type allele with the *glc7-Q48K* allele. The *glc7*Δ could not be examined, since this strain is not viable. We found that the *glc7-Q48K* allele conferred a strong fitness advantage to cells challenged with 2DG, but it came with a detectable fitness cost when cells are grown on glucose. In summary, the relative fitness of the 2DG-resistant alleles could explain why missense alleles were recovered in the *REG1, SNF1* and *GLC7* genes but could not explain why we did not recover any nonsense alleles in *HXK2*.

### Contribution of the Dog2 phosphatase to the 2DG-resistant phenotypes

Overexpression of the DeOxyGlucose (DOG) phosphatases 1 and 2 confers resistance to 2DG (10, 12). We sought to determine what role these phosphatases played in the 2DG resistance caused by the mutations identified in this study. First, we examined the mRNA levels of the *DOG1* and *DOG2* genes using RNAseq. We found that under basal conditions of growth to mid-log phase in synthetic complete media with high glucose, *DOG2* mRNA is expressed at higher levels than *DOG1* mRNA (Fig 7A). Upon shifting cells to low-glucose media for two hours, conditions that activate the Snf1 kinase, we found a significant reduction in mRNA abundance for both *DOG* mRNAs. More germane to this study, both DOG gene mRNAs showed marked increases following a two hour exposure to 2DG. We also examined the abundance of the *DOG* mRNAs in wild type compared to *reg1*Δ and *hxk2*Δ cells. Both deletions confer 2DG resistance, and both lead to a significant increase in *DOG2* but not *DOG1* mRNA abundance (Fig 7B). Increased expression of the *DOG2* mRNA observed in the *reg1*Δ cells was dependent on the Snf1 kinase, since the increase was not observed in the double *reg1Δ snf1*Δ cells. These findings suggested that the underlying mechanism of 2DG resistance was due to increased expression of the Dog2 phosphatase. We tested this idea by measuring 2DG resistance in strains that lacked the *REG1* or *HXK2* genes with or without a deletion covering both the *DOG1* and *DOG2* genes. These genes are adjacent on chromosome 8 and we replaced both genes with the *HIS3* gene to create the double *DOG* deletion (*dog1/2*Δ). As shown previously (5), deletion of *REG1* confers robust resistance to 2DG (Fig 7C). Deletion of both *DOG1* and *DOG2* genes makes cells more sensitive to 2DG, a result recently reported by the Leon lab (12). When *REG1* was deleted from the *dog1/2*Δ strain, the cells were more resistant to 2DG. Thus, deletion of *REG1* was able to confer some 2DG resistance independently of the *DOG1* and *DOG2* genes. An alternative view of this experiment is to note that deletion of the *DOG1* and *DOG2* genes reduced the 2DG resistance of the *reg1*Δ strain. The DOG phosphatases must contribute to some of the 2DG resistance observed in the *reg1*Δ strain. A similar analysis was performed with cells lacking the *HXK2* gene (Fig 7D). Deletion of the *HXK2* gene conferred robust 2DG resistance and further deletion of the *DOG1* and *DOG2* genes only slightly reduced that resistance. Therefore, the DOG phosphatases play a smaller role in the resistance conferred by deletion of *HXK2*. These data suggest that deletion of the *REG1* and *HXK2* genes may confer 2DG resistance by distinct mechanisms and while the DOG phosphatases contribute to some degree to these phenotypes, there must be DOG-independent mechanisms as well.

**Fig. 7.**
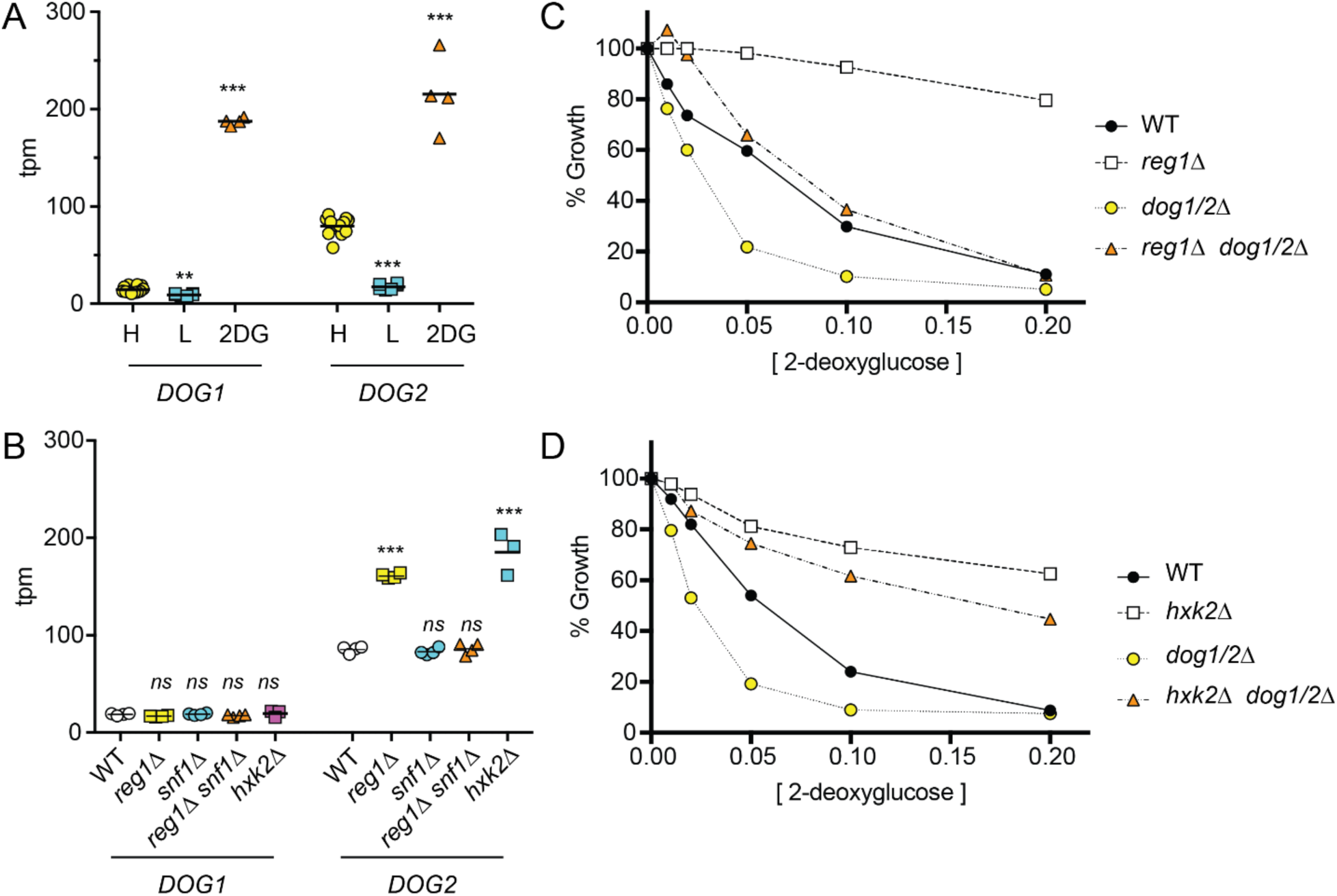
2DG-resistance does not require the DOG phosphatases. **A-B.** Messenger mRNA abundance for the *DOG1* and *DOG2* genes was measured by RNAseq in at least three independent replicates and plotted as transcripts per million mapped reads (tpm). Values statistically different from wild type in high glucose are shown. **A.** RNA was isolated from cells grown in high glucose (H), two hours after shifting to low glucose (L), or two hours after addition of 2DG to a final concentration of 0.1%. **B.** Abundance of *DOG1* and *DOG2* mRNA is shown for wild-type cells, *reg1*Δ, *snf1*Δ, *reg1*Δ *snf1*Δ and *hxk2*Δ cells grown in high glucose. **C and D.** 2DG resistance assays for wild-type (WT) cells and cells with the indicated genotypes.

### Genetic requirements for 2DG resistance conferred by the *SNF1-G53R* allele

In our screen for 2DG-resistant mutants, we uncovered a single dominant allele, *SNF1-G53R.* We took advantage of the dominant inheritance of the 2DG resistance phenotype to test the genetic requirements for 2DG resistance. Our data suggest multiple mechanisms for 2DG resistance act in parallel to confer resistance. Regulation of arrestin-mediated endocytosis of the hexose transporters plays an important role, since deletion of *ROD1* and *ROG3* confers resistance to 2DG (7). Yet the Snf1-G53R kinase conferred additional resistance in the *rod1*Δ *rog3*Δ cells (Fig 8), proving that Rod1- and Rog3-independent mechanisms are also operative. In a similar vein, the DOG phosphatases play an important role in 2DG resistance, since deletion of those genes makes cells more sensitive to 2DG (Figs 7 and 8). Nonetheless, the Snf1-G53R kinase conferred 2DG resistance in the double *dog1/2*Δ delete strain, proving that Snf1 can mediate 2DG resistance via a mechanism that is independent of the DOG phosphatases. We next made strains with deletions in multiple pathways (Rod1 and Rog3 arrestins, Dog phosphatases and Hxk2) to see if we could generate a strain in which the Snf1-G53R kinase could no longer confer resistance. In all combinations of deletions, the Snf1-G53R kinase was able to confer a small but statistically significant level of resistance (Fig 8). Thus, it seems likely that at least one additional Snf1-regulated pathway for 2DG resistance remains to be identified.

**Fig. 8.**
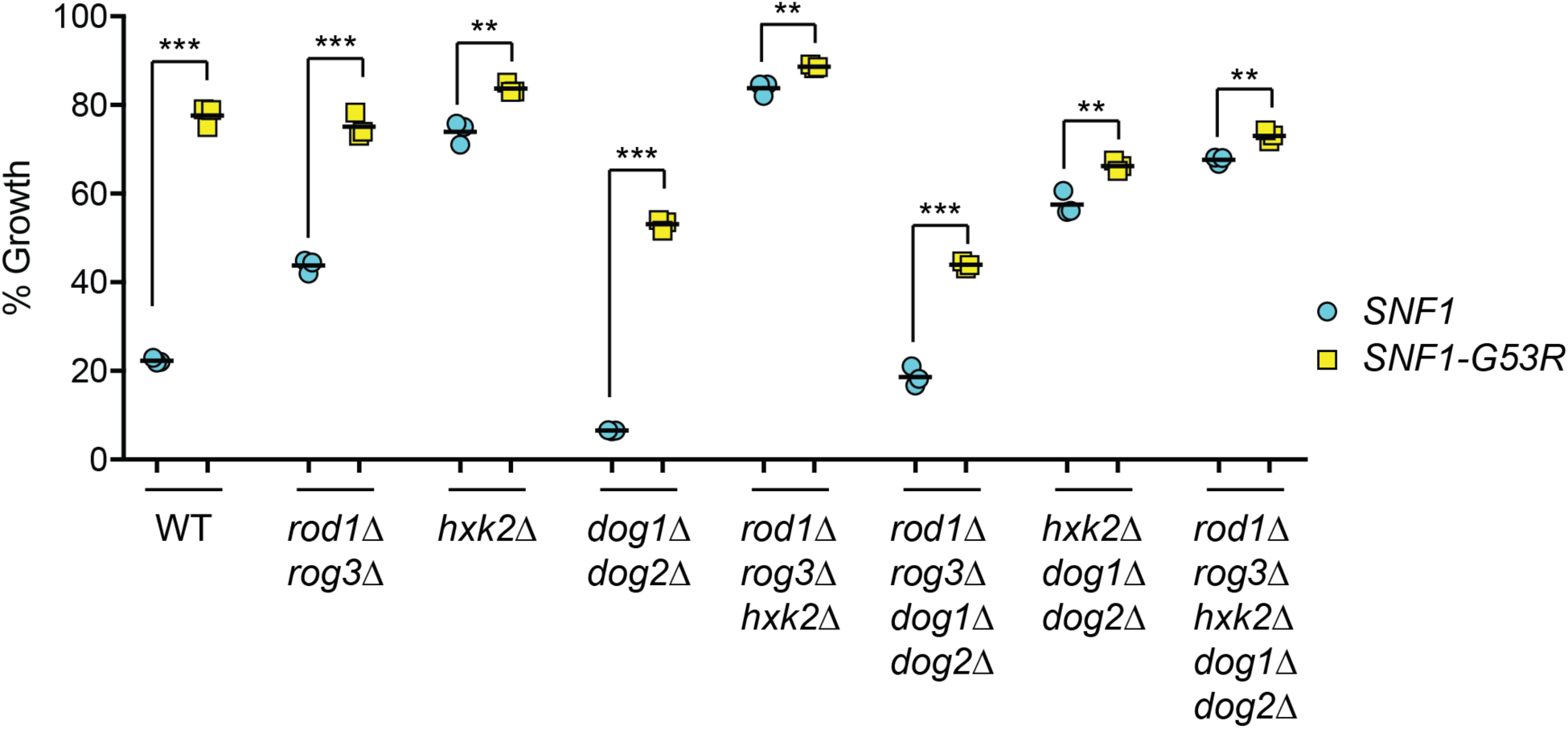
Genetic Requirements for Snf1-G53R-mediated 2DG Resistance. Cells with the indicated genotypes were transformed with low-copy plasmids expressing either wild type Snf1 (blue circles) or Snf1-G53R (yellow squares). Cells were grown overnight in SC media with 2% glucose in the absence or presence of 0.1% 2DG. Growth is expressed as a percentage relative to growth in the absence of 2DG.

## Discussion

The use of 2DG and other metabolic inhibitors as potential cancer therapeutics makes it important to have a deep understanding of their mechanisms of action and the means by which cells acquire resistance. This latter concern is particularly critical for consideration in combinatorial clinical approaches that will block multiple metabolic pathways to more potently inhibit cancer cell progression. In this study, we avoided the use of alternative carbon sources and selected for 2DG resistance in cells growing on glucose as the carbon source, since this would be a growth condition that more closely mimics the plasma glucose used by human tumors. We isolated numerous resistant strains that were in either the hexokinase 2 gene or in genes controlling the Snf1 kinase signaling pathway (*SNF1, REG1,* and *GLC7*).

Yeast express three hexokinase enzymes named Hxk1, Hxk2 and Glk1. All three can phosphorylate glucose, and deletion of all three is needed to significantly block glucose fermentation (Fig S2A). The unanswered puzzle is why *HXK2* mutations are the only hexokinase mutants that confer resistance to 2DG. We can infer that 2-deoxyglucose-6-phosphate (2DG-6P) is the toxic compound, since cells lacking all three hexokinase genes are completely resistant to 2DG (Fig S2B). Both Hxk1 and Hxk2 enzymes can phosphorylate 2DG *in vitro* (Fig S4 and S5), yet only mutations in *HXK2* confer 2DG resistance. While Hxk2 has been described as having both catalytic and gene regulatory activity, we found that our *HXK2* mutants all displayed significant loss of catalytic activity (Fig 1C). In contrast, some alleles showed defects in repression of invertase, notably the *HXK2-G55V* allele, while others (*HXK2-D417G* and *HXK2-R423T*) showed no defect in glucose repression. Mutations in *HXK2* that confer 2DG resistance correlate with loss of catalytic activity, not with defects in *HXK2-*mediated regulation of *SUC2* gene expression (Table 3). The deletion of *HXK2* does cause an increase in the expression of *DOG2* mRNA (Fig 7B) that is not observed in *HXK1* deletion cells (not shown). However, increased expression of *DOG2* mRNA cannot be the entire answer, since deletion of *HXK2* confers 2DG resistance to strains lacking both DOG genes (Fig 7D). A recent study found that mutations affecting all three hexokinase enzymes could reduce glycolytic flux and improve growth in the presence of 2DG (34). Our selection may have identified mutations that reduce the catalytic activity of Hxk2 without inducing expression of Hxk1 thus reducing glycolytic flux. Finally, it is worth noting that our results with various hexokinase mutants differ significantly from those reported in other studies. The N-terminal domain of Hxk2 has been reported to have a Snf1 phosphorylation site at serine 14 that controls nuclear localization (35). Deletion of this N-terminal domain has been reported to deregulate invertase expression (15). We cannot replicate that finding (Fig 1D), even though we are using the same *HXK2* plasmid construct generously provided by the Moreno lab. Our results more closely align with those reported by Botstein’s group, who found that deletion of this region had no effect on invertase expression (17). The only consistent differences in these studies is the use of different *S. cerevisiae* strains: W303 by the Moreno group and S288c by Botstein’s and our group.

Three of the mutations we isolated were in the Snf1 kinase signaling pathway, either in the gene encoding the Snf1 kinase itself or in the PP1 phosphatase subunits that regulate Snf1 activity. The Snf1-G53R allele is a dominant allele first isolated as a partial bypass allele for the deletion of the Snf1 kinase gamma subunit encoded by the *SNF4* gene (14). In an earlier screen in our lab for similar Snf4-independent alleles, we isolated the *SNF1-L183I* allele as a dominant gain-of-function allele that provided significant Snf1 activity in the absence of Snf4 (36). Both Snf1-L183I and Snf1-G53R were isolated in similar screens and both confer resistance to 2DG ((5) and Fig 4A). The L183I residue is in the hydrophobic core of the kinase domain and may stabilize the active confirmation (37). The G53R mutation is on the surface of the kinase domain at the N-terminus, close to the region that interacts with the *β* subunit carbohydrate-binding module (CBM). The interface between the CBM domain and the kinase domain defines an ADaM (allosteric drug and metabolite) site where the direct activators of AMPK bind (38). While both amino acid changes confer a dominant gain-of-function phenotype, it is not clear at a molecular level how they promote 2DG resistance. Neither allele promotes Snf1-mediated phosphorylation of Mig1 (Fig 3D) or of Rod1 (not shown). Similarly, the missense alleles isolated in the PP1 phosphatase enzyme (Reg1-P231L and Glc7-Q48K) do not affect the phosphorylation of Mig1 (Fig 3D), Rod1 (not shown) or the activation loop of Snf1 (Fig 2D and 3B). Thus, the direct target of the Snf1 kinase in the 2DG resistance pathway is not currently known. One possible target for the Snf1 signaling pathway would be a protein that controls the endocytosis of the hexose transporters.

The Snf1 kinase pathway plays an important role in regulating the endocytosis of the hexose (glucose) transporters in both yeast and mammalian cells (7, 9). Yeast cells lacking Snf1 kinase are hypersensitive to 2DG (5) and have increased endocytosis and trafficking of the hexose transporters to the vacuole (7). In this study, we used a strain with an integrated copy of the *HXT3-GFP* gene so that endocytosis could be analyzed directly in our 2DG-resistant strains. Five of the six 2DG-resistant strains we isolated displayed enhanced plasma membrane retention of HXT3-GFP fusion protein (Fig 5). This finding includes two alleles of Hxk2, an enzyme with many reputed activities but not previously linked to protein trafficking.

This study demonstrates that multiple independent and parallel pathways are operating in the resistance to 2DG. Up-regulation of the DOG phosphatases and enhanced plasma membrane retention of the hexose transporters are important determinants of 2DG resistance. Even so, there must be other mechanisms that remain to be discovered. When endocytosis of Hxt3 is diminished by deletion of the *ROD1* and *ROG3* genes (7), we found that Snf1-G53R still promoted additional 2DG resistance (Fig 8). Likewise, when both of the *DOG* phosphatase genes were deleted, Snf1-G53R promoted additional 2DG resistance. Even in strain lacking both arrestins and both DOG phosphatase genes, Snf1-G53R further increased in 2DG resistance. These data suggest that multiple pathways are operating in parallel and that at least one additional pathway remains to be discovered.

## Material and Methods

### Yeast strains and growth conditions

The yeast strains used in this study were all derivatives of S288C. Yeast strains with specific gene deletions were generated in our laboratory or by the *Saccharomyces* Genome Deletion Project (39) and purchased from Thermo Fisher Scientific (Table 4). Cells were grown at 30°C using standard synthetic complete media lacking nutrients needed for plasmid selection (40).

### Selection of spontaneous 2DG-resistant mutants

Spontaneous mutations that conferred 2DG resistance were selected in the haploid strain MSY1333 (Table 1). Approximately 2×10^7^ cells were spread on agar plates containing synthetic complete media with 2% glucose (g/100ml) and 0.1% 2-deoxyglucose. Plates were incubated at 30°C for 4-6 days, and colonies were isolated for further study.

### Whole Genome Sequencing analysis of 2DG-resistant mutants

Spontaneously 2DG-resistant strains were analyzed for mutations by whole genomic sequencing using pooled linkage analysis (41) when possible. Genomic DNA was extracted using a glass bead phenol extraction method (42). Sequencing libraries were prepared using a modified Illumina Nextera protocol and multiplexed onto a single run on an Illumina NextSeq500 to produce 151-bp paired-end reads (13). Sequencing produced an average depth of 10-20 million reads per sample. Reads were mapped against the reference genome of strain S288C and variants detected using CLC Genomics Workbench (Qiagen).

### Mutagenesis and Epitope Tagging of plasmid constructs

Oligonucleotide-directed mutagenesis was performed with Pfu polymerase, followed by digestion of the starting plasmid template with the restriction enzyme *DpnI* (43). All mutations reported in the study were confirmed by DNA sequencing. The wild-type *REG1* and *HXK2* alleles as well as mutant alleles were modified to contain three copies of V5 epitope (44). The Snf1 and Glc7 proteins were epitope tagged with three tandem copies of the HA epitope placed at the C-terminus of the open reading frame (20, 25). All of these constructs were introduced to yeast cells on low-copy CEN plasmids (45) and were expressed from their respective native promoters. To create a yeast strain expressing HA-tagged Glc7-Q48K, a diploid yeast strain heterozygous at the *GLC7* locus (*GLC7/glc7Δ::KAN)* was transformed with *GLC7* plasmids under *LEU2* selection. After sporulation, haploid segregants that were KAN+ and LEU+ were collected. These haploid cells contained the *glc7Δ::KAN* allele on chromosome five but were viable due to the presence of HA-tagged *GLC7* or *glc7-Q48K* on a plasmid.

### 2-deoxyglucose resistance assays

2DG resistance was measured in liquid cultures as previously described (5). Briefly, overnight cultures were diluted to A_600_ of 0.1 and grown in the absence of 2DG or in the presence of increasing concentrations of 2DG (0.01%, 0.02%, 0.05%, 0.1%, 0.2%). Cells were grown for 18hr at 30°C. Each A_600_ was measured, and cell growth was normalized to growth in the absence of 2DG for each strain.

### Enzyme assays

The invertase activity of log-phase cells grown in high glucose or 2 hours after shifting to low glucose media was quantitatively assayed used a colorimetric assay coupled to glucose oxidase (46). Three independent cultures were assayed, and the mean value is plotted with standard error indicated. The units of invertase activity used were mU/OD, where 1 U equals one μmole glucose released per minute. Hexokinase enzyme activity was measured by coupling the phosphorylation of glucose to its oxidation by glucose-6-phosphate dehydrogenase (17, 18). Assays were conducted in 100 µl reactions containing 50 mM Tris-Cl pH 7.4, 10 mM MgCl_2_, 5% glycerol (vol/vol) 1 mM ATP, 10 mM glucose, 0.3 U glucose-6-phosphate dehydrogenase (Sigma G7877) and 0.1 mM NADP^+^. Production of NADPH was determined in a 96-well plate reader by measuring absorbance at 340 nm using the extinction coefficient of 6,220 M^-1^ cm^-1^. Units of hexokinase activity were expressed as nmoles/min. Hexokinase enzymes were assayed in yeast whole cell extracts prepared from cells lacking all three yeast hexokinase genes (MSY1477) and expressing a single hexokinase from a low-copy plasmid.

### Protein extractions

Trichloroacetic acid (TCA) was utilized for extraction of the Reg1 using a modification of a method previously reported (47). Log-phase cells (2.5 OD’s) were harvested and washed with cold water before being resuspended in 1 ml of cold sterile water. Next, 150 ul of 2N NaOH and 78.3 ul of 1.12 M β-mercaptoethanol were added, and samples incubated on ice for 15 minutes. To precipitate proteins, 150 ul of cold TCA (50%) was subsequently added and incubated for 20 more minutes on ice. Samples were centrifuged at 4°C for 5 minutes, and pellets were resuspended in 50 ul TCA sample buffer (80 mM Tris-Cl pH 8.0, 8 mM EDTA, 120 mM DTT, 3.5% SDS, 0.29% glycerol, 0.08% Tris base, 0.01% bromophenol blue). Samples were incubated at 37°C for 30 minutes, and cell debris was removed by centrifugation prior to gel electrophoresis.

Glass bead extracts were prepared from liquid cultures grown to mid-log (OD_600_ of 0.6 to 0.8). 25 mL of cells were collected by centrifugation at 4°C at 3500 rpm and washed once in extraction buffer (20 mM Hepes pH 7, 0.5 mM EDTA, 0.5 mM DTT, 5 mM MgAc, 0.1 M NaCl, 10% glycerol). The cell pellet was resuspended in extraction buffer (4 packed cell volumes) supplemented with added protease inhibitors. Glass beads equal to the volume of the pellet were added, and the cells were shaken in an MP FastPrep-24 for three cycles of 20 seconds with 5 minutes of rest on ice between each cycle. Cell debris was removed by centrifugation at maximum speed for 5 minutes. Extracts were frozen and stored at −80°C.

### Western blotting

Western blotting techniques were modified from previous methods (25, 48). Proteins tagged with the HA epitope (Snf1, Glc7) were detected with Anti-HA probe (Santa Cruz) diluted 1:2,000. Goat anti-mouse IRDye 800CW (Li-Cor) diluted 1:5,000 was used as the secondary antibody. For detection of phosphorylated Snf1, Phospho-AMPKalpha (Thr172) antibody (Cell Signaling) diluted 1:1,000 was used. Goat anti-rabbit IRDye 800CW (Li-Cor) was used as the secondary antibody at a 1:10,000 dilution. Proteins tagged with the V5 epitope (Hxk2, Reg1) were detected with the Anti-V5 probe (Invitrogen) diluted 1:1,000. Goat anti-mouse IRDye 680 (Thermo) diluted 1:10,000 was used as the secondary antibody. Blots were visualized using an Odyssey Infrared Imager (Li-Cor), and band quantification was performed using Odyssey software.

### Two-hybrid analysis

Two-hybrid interactions were analyzed as previously described (49). Interactions between proteins were assessed by growth on medium containing 2% glucose lacking histidine, using strains with the *GAL1-HIS3* reporter integrated at the *LYS2* locus. The positive control plasmids used in this study encode the herpes simplex virus capsid protein 22a (50). The entire open reading frame for the proteins Snf1 and Glc7 and their derivatives Snf1-G53R and Glc7-Q48K were expressed as a fusion to the Gal4 DNA binding domain in pGBT9 (51). The entire open reading frame for the protein Reg1 and its derivative Reg1-P231L was expressed as a fusion to the Gal4 activation domain in the plasmid pACT2 (Clontech).

### RNAseq analysis

RNA samples were prepared from multiple independent yeast cultures grown on synthetic complete medium using the RNeasy Mini Kit (Qiagen). Sequencing libraries were prepared using the TruSeq Stranded mRNA library method (Illumina). RNA sequences were mapped to *S. cerevisiae* mRNA using the kallisto software package (29). Each RNA sample yielded 40-50 million reads. mRNA abundance was expressed in transcripts per million mapped reads (tpm). Complete datasets for the RNAseq experiments described here are presented in the attached file as Table S5.

### Fluorescence microscopy and quantification

Cells were grown to mid-log phase in synthetic complete medium containing 2% glucose and then incubated an additional 2 h after 0.2% glucose or 0.2% 2DG addition. Fluorescent images were acquired on a TiE2 inverted microscope (Nikon, Chiyoda, Tokyo, Japan) equipped with an Apo100x objective (NA 1.45) and captured with an Orca Flash 4.0 cMOS (Hammamatsu, Bridgewater, NJ) camera and NIS-Elements software (Nikon). To define vacuoles, cells were incubated with 250 uM Cell Tracker Blue CMAC dye (Life Technologies, Carlsbad, CA). All images were acquired using equivalent parameters and are presented as evenly adjusted images from Adobe Photoshop. For all figures an unsharp mask was applied with a threshold of 5 and a pixel radius of 5. Quantification of plasma membrane and vacuolar fluorescence intensities was assessed as described (7).

### Statistical significance

Unless otherwise stated, mean values represents the average for a minimum of three independent measurements, and the error bars represent 1 standard error. Statistical significance was determined using the Student t test for unpaired variables with equal variance. For statistical analysis of HXT3-GFP localization, populations were assessed for statistically significant differences using non-parametic Kruskal-Wallis statistical analyses with Dunn’s multiple comparisons post-hoc test using Prism software (GraphPad, La Jolla, CA). In all cases, p values are indicated as follows: * p<0.05; ** p<0.01; *** p<0.001.

## Acknowledgements

This research was funded by the National Science Foundation (MCB CAREER 1902859 to A.F.O.) and the National Institutes of Health (R01 GM46443 to M.C.S.). We thank Dan Snyder and Vaughn Cooper for whole genome sequencing, the University of Pittsburgh Genomics Research Core for RNA sequencing, Mitchell Ellison for generously sharing his bioinformatics skills, Sebastien Leon for sharing results prior to publication and Fernando Moreno for the gift of *HXK2* plasmids.

